# Reimplementing Unirep in JAX

**DOI:** 10.1101/2020.05.11.088344

**Authors:** Eric J. Ma, Arkadij Kummer

**Affiliations:** Scientific Data Analysis, NIBR Informatics, Novartis Institutes for Biomedical Research; Bioreactions Group, Global Discovery Chemistry, Novartis Institutes for Biomedical Research

## Abstract

UniRep is a recurrent neural network model trained on 24 million protein sequences, and has shown utility in protein engineering. The original model, however, has rough spots in its implementation, and a convenient API is not available for certain tasks. To rectify this, we reimplemented the model in JAX/NumPy, achieving near-100X speedups in forward pass performance, and implemented a convenient API for specialized tasks. In this article, we wish to document our model reimplementation process with the goal of educating others interested in learning how to dissect a deep learning model, and engineer it for robustness and ease of use.

## Introduction

UniRep is a recurrent neural network, trained using self-supervision on 24 million protein sequences to predict the next amino acid in a sequence (Alley et al. 2019). Its most powerful model allows for embedding arbitrary length sequences in a 1900-long feature vector that can be used as the input to a “top model” for unsupervised clustering or supervised prediction of protein properties.

The original model was implemented in TensorFlow 1.13 (Abadi et al. 2016), and its original API only allowed for one sequence to be transformed at once. While test-driving the model, we observed two problems with it. The first is that that the original implementation took an abnormally long amount of time to process multiple sequences, requiring on the order of dozens of seconds to process single sequences. The second was that its API was not sufficiently flexible to handle multiple sequences passed in at once; to get reps of multiple sequences, one needed to write a manual for-loop, re-using a function inside which returns the reps for a single sequence. When fine-tuning model weights, sequences needed to be batched and padded to equal lengths before being able to be passed in to the model. Neither appeared to be user-friendly.

Thus, while the model itself holds great potential for the protein engineering field, the API prevents us from using it conveniently and productively. We thus sought to reimplement and package the model in a way that brings a robust yet easy-to-use experience to protein modellers and engineers.

In particular, our engineering goals were to provide:

- A function that can process multiple sequences of arbitrary lengths,
- Vectorizing the inputs to make it fast.
- A single function call to “evotune” the global weights.

## Profiling tf-unirep and jax-unirep

To investigate the performance of the original and our reimplementation, we used Python’s `cProfile` facility to identify where the majority of time was spent in the respective codebases. The functions used for profiling were:

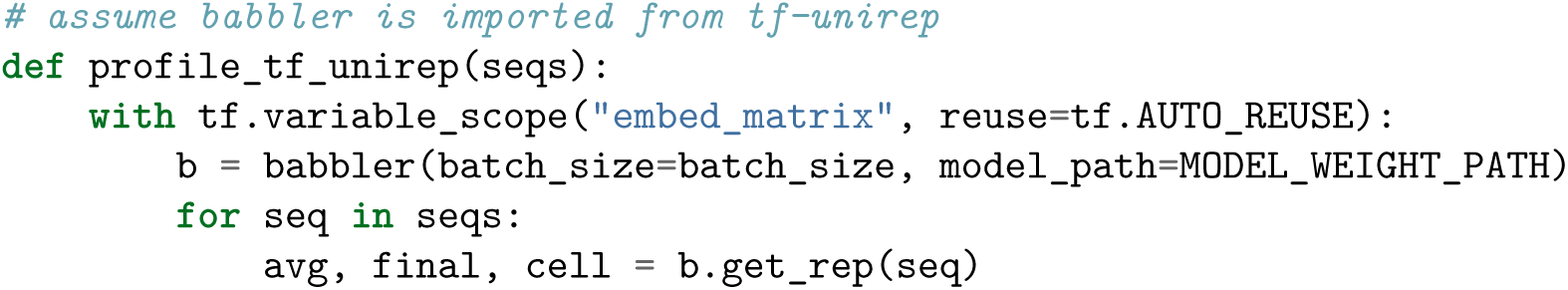

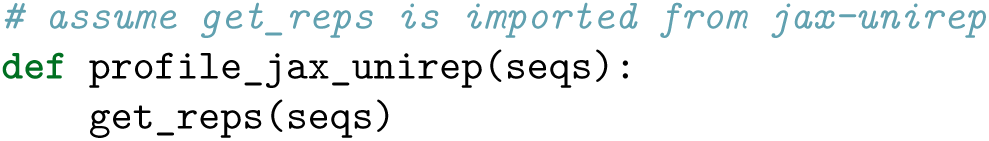

We then used SnakeViz to visualize the code execution profile results.

As is visible from the code execution flamegraph, the unreasonably long time that it takes to process ten sequences was probably due to the time spent in TensorFlow’s session. Because of TensorFlow’s compiled nature, we thus deduced that the majority of the execution time was most likely in the graph compilation phase. Unfortunately, cProfile could not give us any further detail beyond the `_pywrap_tensorflow_internal.TF_SessionRun_wrapper` in the call graph, meaning we were unable to conveniently peer into the internals of TF execution without digging further.

On the basis of this profiling, we hypothesized that the cause of speed problems was graph compilation in TF1.x, and that we could obtain speedups by using a non-graph-compiled tensor library. There were three choices for us at this point: TF2.x, PyTorch and JAX, and we chose the latter. Our choice was motivated by the following reasons:

1. JAX uses the NumPy API, which is idiomatic in the Python scientific computing community.
2. JAX provides automatic differentiation, which would enable us to reimplement weights fine-tuning.
3. JAX encourages functional programming, which makes implementation of neural network layers different from class-based implementations (e.g. PyTorch and Keras). This was an intellectual curiosity point for us.
4. JAX is “eagerly” executable like PyTorch and TF2, which aids debugging.

Besides these *a priori* motivating reasons, we also uncovered other reasons to use JAX midway:

1. JAX’s compiled and automatically differentiable primitives (e.g. `lax.scan`) allowed us to write performant RNN code.
2. `jit` and `vmap` helped with writing performant training loops.

We thus reimplemented the model in JAX/NumPy. (See “Reimplementation Main Points” section for details.) As is visible from the code profiling APIs that we used, we designed a cleaner and more expressive API that could be faster and handle multiple sequences of variable lengths, without introducing the mental overhead of TensorFlow’s complex scoping syntax. An expressive and clean API was something that we would expect a computational protein engineer would desire, as having this API form would lower mental overhead while also hopefully being faster to execute and write.

A formal speed comparison using the same CPU is available below.

We also needed to check that our reimplementation correctly embeds sequences. To do so, we ran a dummy sequence through the original and through our reimplementation, and compared the computed representations. Because it is 1900-long, a visual check for correctness is a trace of 1900-long embedding.

We also verified that the embeddings calculated using the pre-trained weights were informative for top models, and trained a model to predict the brightness of around 50’000 avGFP variants (as the authors did). avGFP is a green-fluorescent protein that has been extensively studied in the literature. Many studies generated mutants of this protein, measuring the changes in brightness for each mutant, to try to understand how protein sequence links to function or simply to increase brightness.

We binarized brightness values into a “dark” and a “bright” class, and used scikit-learn’s implementation of logistic regression for classification. Average performance across 5-fold cross-validation is shown in Figure 3. (avGFP data came from (Sarkisyan et al. 2016).)

**Figure 1:**
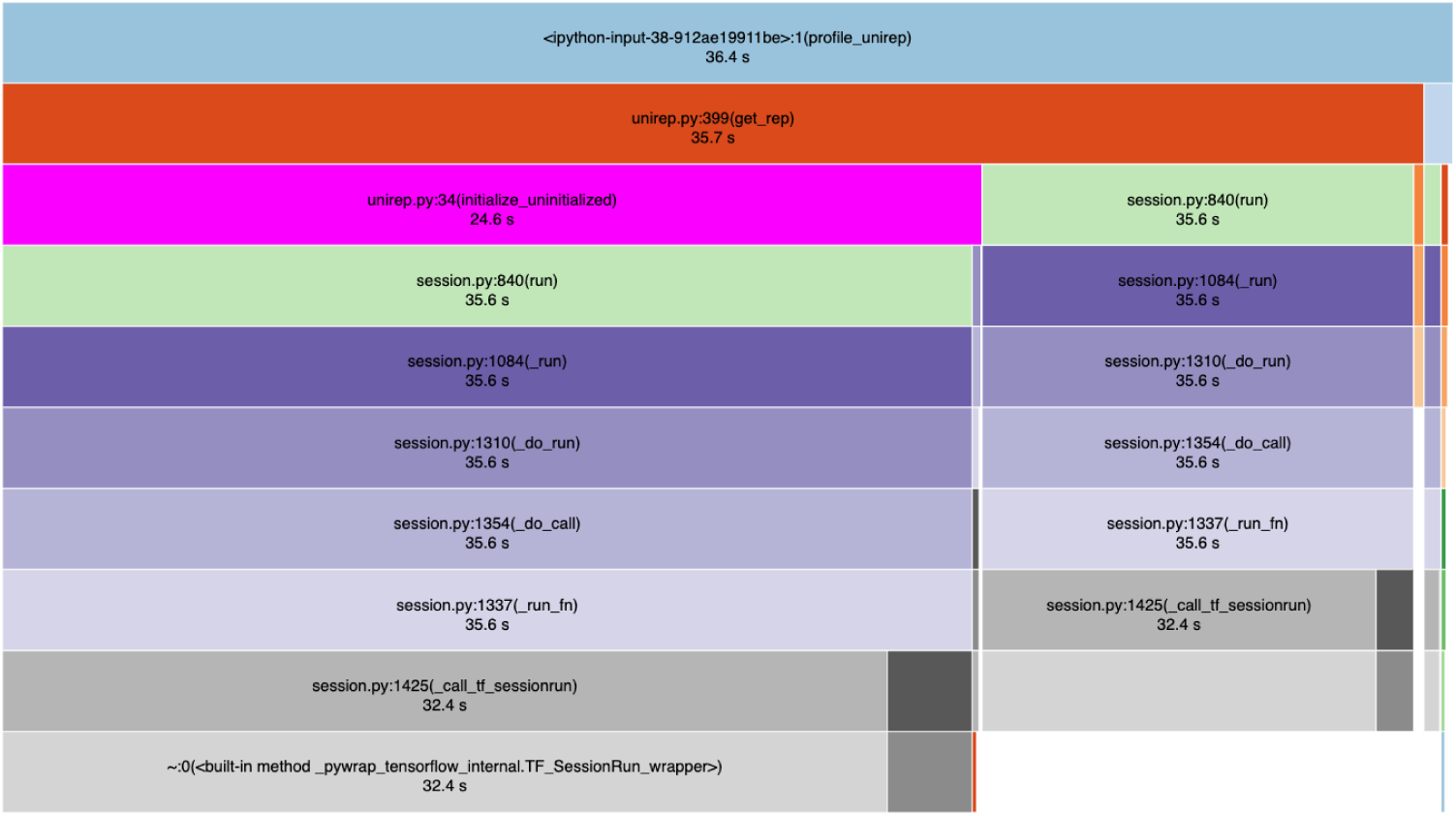
Flame graph of the original UniRep’s implementation, down to 10 levels deep from the profiling function that was called.

**Figure 2:**
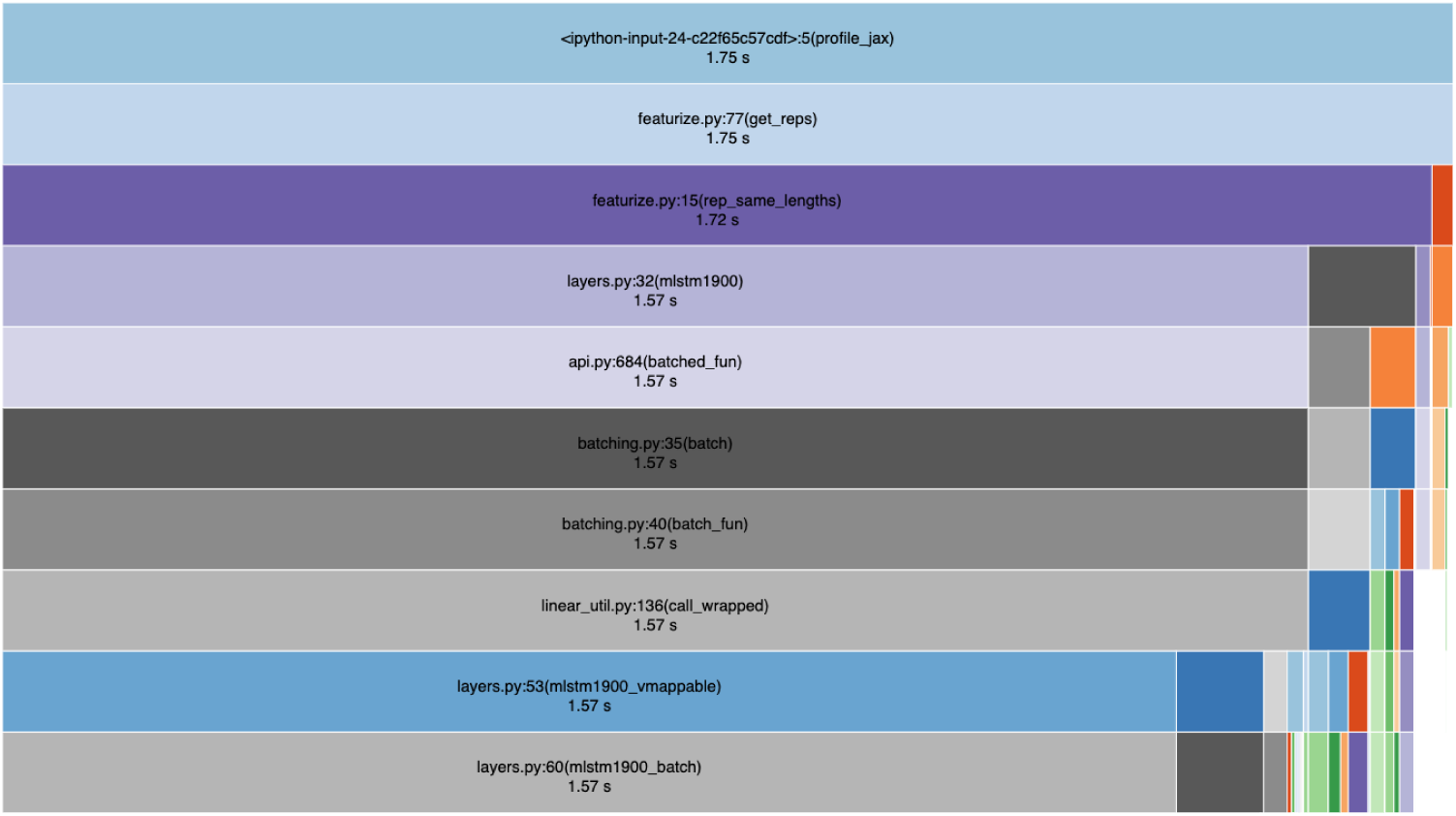
Flame graph of the jax-unirep reimplementation, down to 10 levels deep from the profiling function that was called.

**Figure 3:**
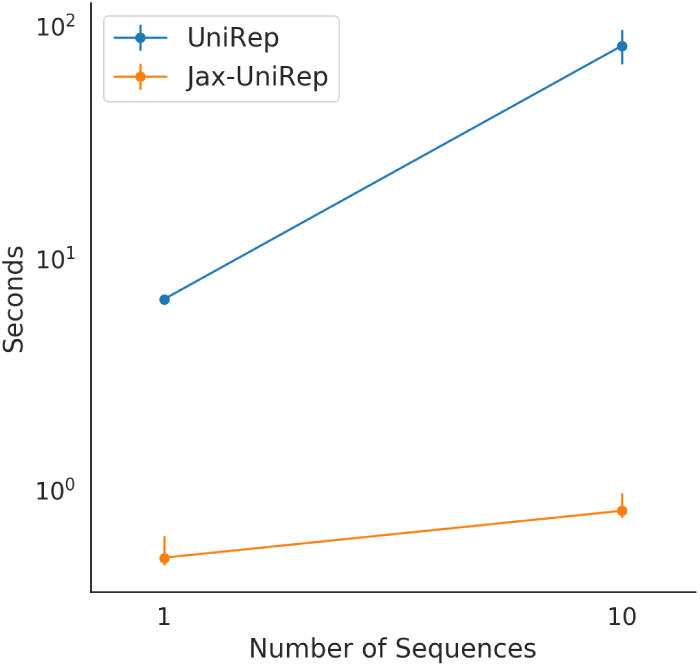
Speed comparison between the original implementation (UniRep) and our re-implementation (Jax-UniRep). Both one and ten random sequences of length ten were transformed by both implementations. Our re-implementation could make use of vectorization in the multi-sequence case, whereas in the original implementation the sequences were transformed one at a time.

**Figure 4:**
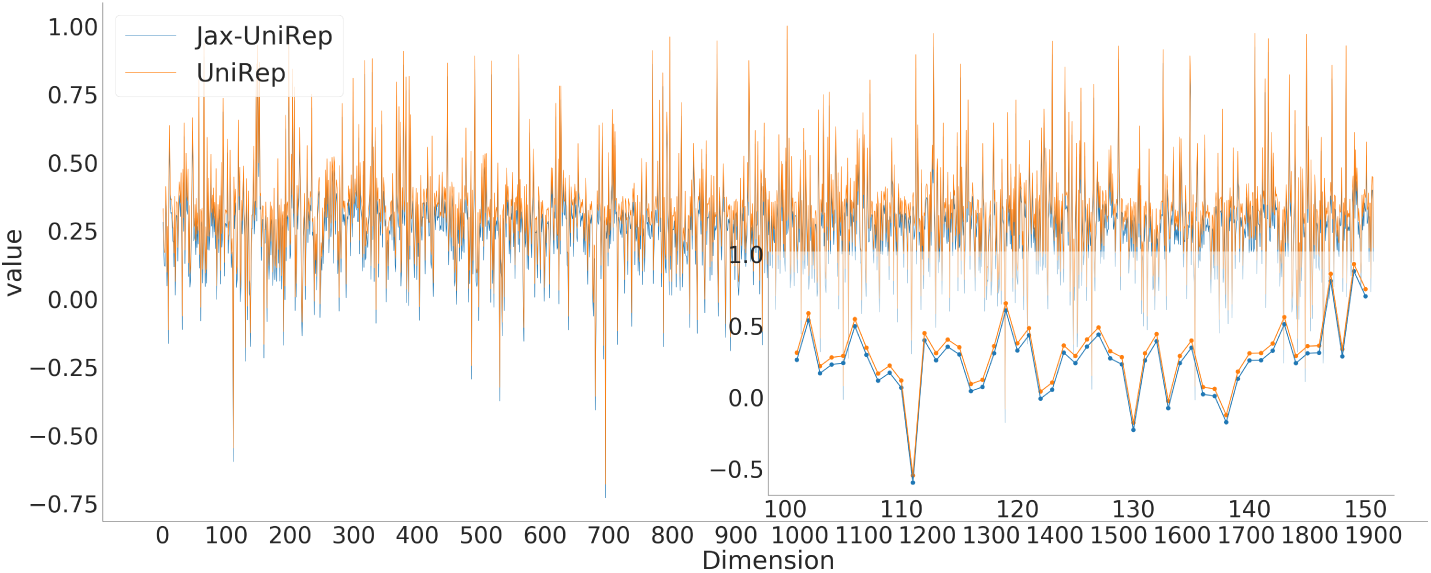
Comparison of the average hidden state between the implementations when transforming the same sequence. Because the two traces of the hidden state dimensions overlapped almost perfectly, a small constant was added to the UniRep values, such that both traces become visible. The inset shows 50 out of the total 1900 dimensions.

**Figure 5:**
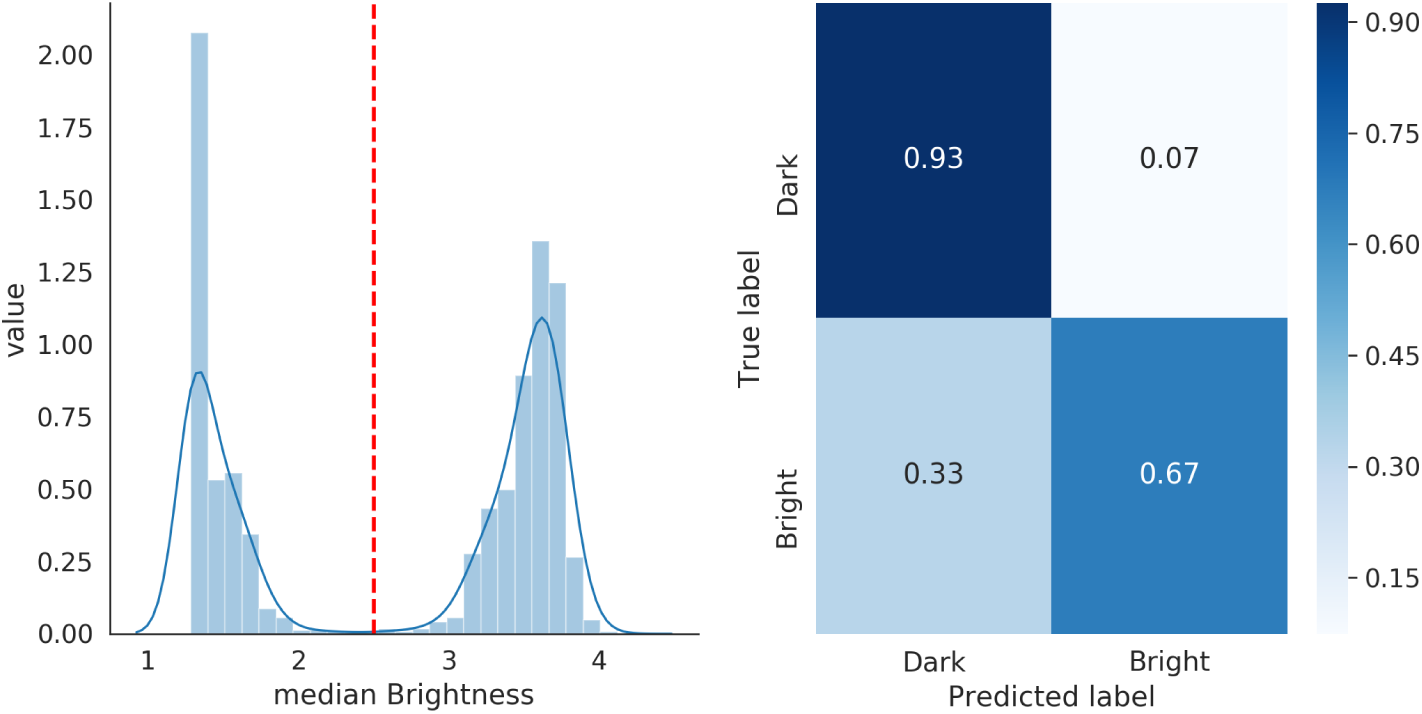
GFP brightness classification using a logistic regression top model taking in the 1900-long average hidden state representations of the GFP protein sequences. Left: Distribution of GFP brightness values in the dataset. Red dotted line indicates classification breakpoint. Points to the left get labeled as “Dark”, while points to the right get labeled “Bright”. Right: Confusion matrix showing the classification accuracy of the model.

## Reimplementation Main Points

### Choice of JAX

JAX was our library choice to reimplement it in, because it provides automatic differentiation machinery (Bradbury et al. 2018) on top of the highly idiomatic and widely-used NumPy API (Oliphant 2006). JAX uses a number of components shared with TensorFlow, in particular the use of the XLA (Accelerated Linear Algebra) library to provide automatic compilation from the NumPy API to GPU and TPU.

Part of the exercise was also pedagogical: by reimplementing the model in a pure NumPy API, we are forced to become familiar with the mechanics of the model, and learn the translation between NumPy and TensorFlow operations. This helps us be flexible in moving between frameworks.

Because JAX provides automatic differentiation and a number of optimization routines as utility functions, we are thus not prohibited from fine-tuning UniRep weights through gradient descent.

During the reimplementation, we also discovered that JAX provided convenient utilities (`lax.scan`, `vmap`, and `jit`) to convert loops into fast, vectorized operations on tensors. This had a pleasant effect of helping us write more performant code. We were also forced to reason clearly about the semantic meaning of our tensor dimensions, to make sure that vecotrization happened over the correct axes. We commented at every tensor operation step how the shapes of our input(s) and output(s) should look like. One example from our source:

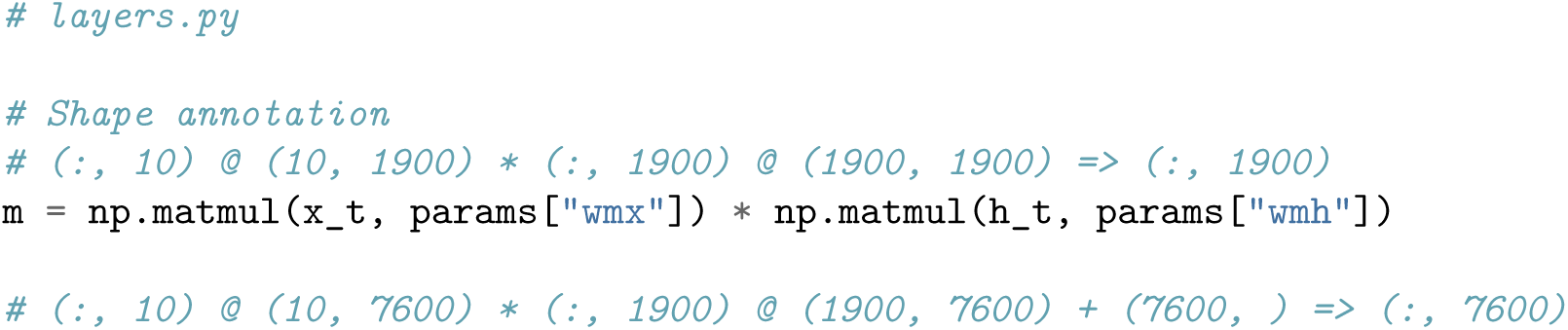

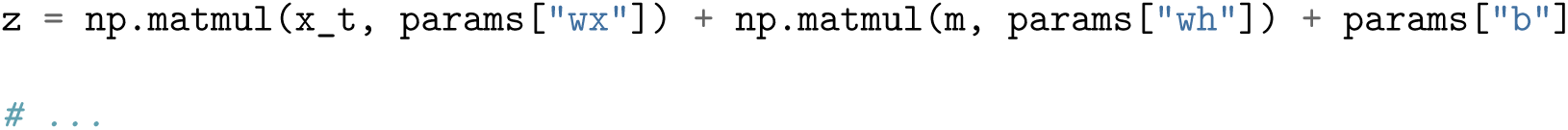

### Tensor Ops Reimplementation

The process of tensor ops reimplementation were as follows.

Firstly, we started from the RNN cell (`mLSTM1900_step`), which sequentially walks down the protein sequence and generates the single step embedding. We thus end up with a “unit cell” function:

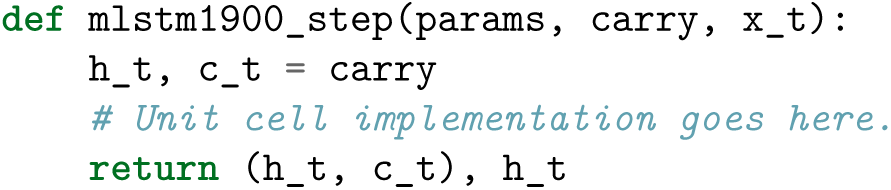

Secondly, we wrapped the RNN cell using `lax.scan` to scan over a single sequence. This is the `mlstm1900_batch` function:

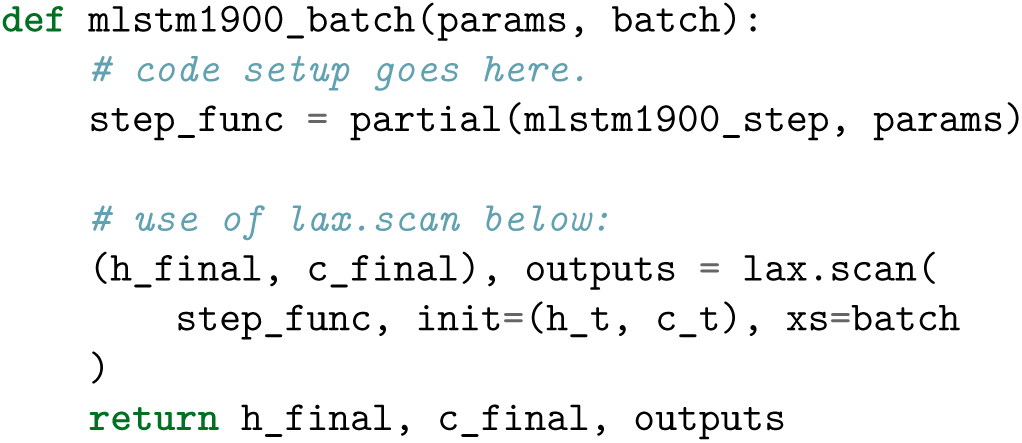

Thirdly, we then used `jax.vmap` to vectorize the operation over multiple sequences, thus generating `mlstm1900`:

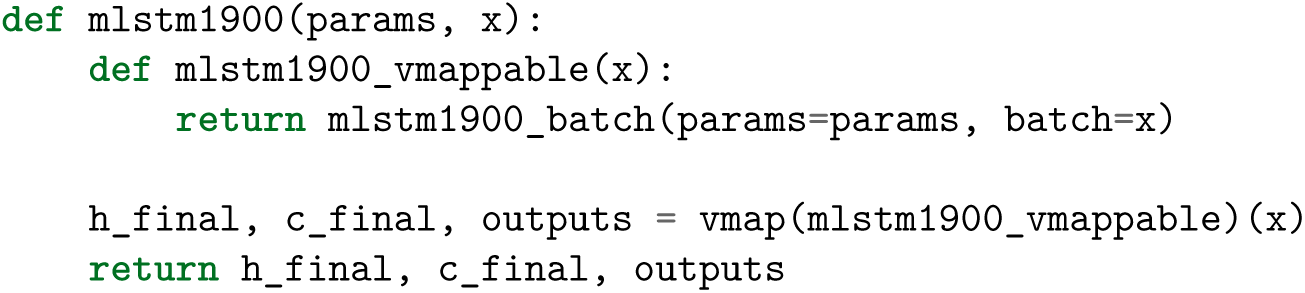

Effectively, `jax.vmap` and `lax.scan` replace for-loops that we would otherwise write, which would incur Python type-checking overhead that would accumulate. `lax.scan` being effectively a pre-compiled for-loop enables pre-allocation of the necessary memory needed for backpropagation, which also contributes to a speedup. As the for-loop type checking penalty is well-known in Python, a detailed comparison between `jax.vmap`, `lax.scan`, and a vanilla `for` loop is out of scope for this paper. The full source code is available in `jax_unirep/layers.py`.

Besides reimplementation, we also took care to document the semantic meaning of tensor dimensions. This had the pleasant side effect of forcing us to order our tensor dimensions in a sane fashion, such that the “batch” or “sample” dimension was always the first one, with explicit documentation written to guide a new user on this convention.

While reimplementing the model, we also generated a test suite for it. Most of our tests check that the shapes of returned tensors were correct. For the unit RNN cell, we provided an example-based test with random matrices. The same applied to the batch function. However, for the full forward model, we provided a property-based test, which checked that tensor dimensions were correct given different numbers of samples. These are available in the source `tests/` directory. As a known benefit with software testing, our tests allowed us to rebuild the full model piece by piece, while always making sure that each new piece did not break the existing pieces.

### Utility Reimplementation

For the `get_reps()` functionality, we copied quite a bit of source code from the original, including the original authors’ implementation of embedding a sequence into an *l*-by-10 embedding matrix first. However, we added tests to guarantee that they were robust, as well as technical documentation to clarify how it works.

We did this because one way that deep learning models can be fragile is that the input tensors can be generated incorrectly but still have the expected shapes. Thus, though the structure of input tensors might be correct, their semantic meaning would be completely wrong. (Adversarial examples can be generated this way.) Thus, the input to the model has to be carefully controlled. Moreover, input tensors are *not* the raw-est form of data; for a protein engineer, the protein sequence is. Thus, having robustly tested functions that generate the input tensors with correct semantic meaning is crucial to having confidence that the model works correctly end-to-end.

### APIs

Because we expect the model to be used as a Python library, the model source and weights are packaged together. This makes it much more convenient for end-users, as the cognitive load of downloading starter weights is eliminated.

The `get_reps()` function is designed such that it is flexible enough to accept a single sequence or an iterable of sequences. This also reduces cognitive load for end-users, some of whom might want to process only a single sequence, while others might be operating in batch mode. `get_reps()` also correctly handles sequences of multiple lengths, further simplifying usage for end-users. In particular, we spent time ensuring that `get_reps()` correctly batches sequences of the same size together before calculating their reps, while returning the reps in the same order as the sequences passed in. As usual, tests are provided, bringing the same degree of confidence as we would expect from tested software.

## Lessons Learned

We found the reimplementation exercise to be highly educational. In particular, we gained a mechanical understanding of the model, and through documenting the model functions thoroughly with the semantic meaning of tensor dimensions, we were able to greatly reduce our own confusion when debugging why the model would fail.

During the reimplementation, we found the “sigmoid” function to be an over-loaded term. We initially used a sigmoid that had an incorrect slope, yielding incorrect reps. Switching to the correct sigmoid slope rectified the problem. A similar lesson was learned while reimplementing the L2 norm of our weights.

Writing automated tests for the model functions, in basically the same way as we would test software, gave us the confidence that our code changes would not inadvertently break existing functionality that was also already tested. We also could then more easily narrow down where failures were happening when developing new code that interacted with the model (such as providing input tensors).

Through reimplementation, we took the opportunity to document the semantic meaning of tensor axes and their order, thus enabling ourselves to better understand the model’s semantic structure, while also enabling others to more easily participate in the model’s improvement and development.

Competing tensor libraries that do not interoperate seamlessly means data scientists are forced to learn one (and be mentally locked in). To break free of framework lock-in, being able to translate between frameworks is highly valuable. Model reimplementation was highly beneficial for this.

UniRep was implemented in Tensorflow 1.13. It is well-known that TF1’s computation graph-oriented API does not promote ease of debugging in native Python. Hence, it may sometimes be difficult to find spots in a TF model where one could speed up computations. By instead treating neural network layers as functions that are eagerly evaluated, we could more easily debug model problems, in particular, the pernicious tensor shape issues.

We believe that the speedup that we observed by reimplementing in JAX came primarily from eliminating graph compilation overhead and an enhanced version of the original API design. In anecdotal tests, graph compilation would take on the order of seconds before any computation occurred. Because the original implementation’s `get_reps` function did not accept multiple sequences, one had to use a for-loop to pass sequence strings through the model. If a user were not careful, in a worst-case scenario, they would end up essentially paying the compilation penalty on every loop iteration.

By preprocessing strings in batches of the same size, and by keeping track of the original ordering, then we could (1) avoid compilation penalty, and (2) vectorize much of the tensor operations over the sample axis, before returning the representation vectors in the original order of the sequences. In ensuring that the enhanced `get_reps` API accepted multiple sequences, we also reduced cognitive load for a Python-speaking protein data scientist who might be seeking to use the model, as the function safely handles a single string and an iterable of strings.

An overarching lesson we derive from this experience is as follows. If “models are software 2.0” (Karpathy 2017), then data science teams might do well to treat fitted model weights as software artefacts that are shipped to end-users, and take care to design sane APIs that enable other developers to use it in ways that minimize cognitive load.

## Future Work

As we have, at this point, only implemented the 1900-cell model. Going forth, we aim to work on implementing the 256- and 64-cell model.

Evotuning is an important task when using UniRep (Alley et al. 2019), and we aim to provide a convenient API through the `evotune()` function. Here, we plan to use Optuna to automatically find the right hyperparameters for finetuning weights, using the protocol that the original authors describe. This would enable end-users to “set and forget” the model fitting protocol rather than needing to babysit a deep learning optimization routine. Like `get_reps()`, `evotune()` and its associated utility functions will have at least an example-based test, if not also a property-based test associated with them.

Community contributions and enhancements are welcome as well.

## Software Repository

`jax-unirep` is available on GitHub at https://github.com/ElArkk/jax-unirep.

## Acknowledgments

We thank the UniRep authors for open sourcing their model. It is our hope that our reimplementation helps with adoption of the model in a variety of settings, and increases its impact.

